# Polygenic risk score for schizophrenia is not strongly associated with the expression of specific genes or gene sets

**DOI:** 10.1101/205518

**Authors:** David Curtis

**Affiliations:** UCL Genetics Institute, UCL, Darwin Building, Gower Street London WC1E 6BT.; Centre for Psychiatry, Barts and the London School of Medicine and Dentistry, Charterhouse Square, London EC1M 6BQ

**Keywords:** Schizophrenia, polygenic risk score, RNA, expression

## Abstract

The polygenic risk score (PRS) is derived from SNPs including both those which are genome-wide significant and also a large number of others more weakly associated with schizophrenia. Such variants are widely dispersed, though concentrated near genes expressed in the brain, and it has been proposed that these SNP associations result from impacts on cell regulatory networks which ultimately affect the expression or function of a modest number of "core" genes. A previous study demonstrated association of some GWAS-significant variants with expression of a number of genes, by examining pair-wise correlations of gene expression with SNP genotypes. The present study used data downloaded from the CommonMind Consortium site, consisting of SNP genotypes and RNAseq expression data from the dorsolateral prefrontal cortex, to examine whether the expression of individual genes or sets of genes correlated with PRS in 207 controls and 209 schizophrenia cases. Although the PRS was significantly associated with phenotype, the correlations with genes and genes sets followed distributions expected by chance. Thus, this analysis failed to demonstrate that the PRS captures a cumulative effect of multiple variants impacting the expression of a small number of genes and it failed to focus attention on a small number of genes of biological relevance. The multiple SNP associations observed in schizophrenia may result from other mechanisms, including effects mediated indirectly through environmental risk factors.

## Introduction

There is a substantial genetic contribution to the aetiology of schizophrenia which in part manifests as association with common variants, with over 100 loci meeting conventional standards for significance in a genome-wide association study (Schizophrenia Working Group of the Psychiatric Genomics Consortium, 2014). The same study showed that a polygenic risk score (PRS) was more strongly predictive of schizophrenia if it included not only the variants which were statistically significant but also many other variants with only weak evidence for association. This clearly demonstrated that there are a very large number of common variants whose frequency differs between cases and non-cases. In a recent review of this field, the authors noted the prior evidence for the involvement of large numbers of variants which were very widely distributed across the genome and proposed that the term "omnigenic" could appropriately be used (Boyle et al., 2017). However in their own analyses they demonstrated that only variants near genes expressed in the brain contribute substantially to heritability of schizophrenia and that variants near genes expressed specifically in the brain made a larger contribution per variant than those near more widely expressed genes. They also noted that common variants show only a modest enrichment for genes by functional category and contrasted this with the findings from studies of rare variants and CNVs which implicate synaptic and neuronal genes (Fromer et al., 2014; Marshall et al., 2016; Purcell et al., 2014). Although they did not refer to it, an additional study of ultra-rare, gene disruptive variants also demonstrated enrichment in these categories (Genovese et al., 2016). To make sense of these observations, they proposed that a given phenotype might be directly affected by a modest number of "core" genes but that, because cell regulatory networks are highly interconnected, any expressed gene is likely to have some effect on the regulation or function of these core genes.

A previous study sought to investigate the relationship between common variants and gene expression by examining SNP genotypes and RNAseq results from the dorsolateral prefrontal cortex (DLPFC) (Fromer et al., 2016). This reported that about 20% of schizophrenia-associated loci contained variants which could contribute to altered gene expression or liability. Considering SNP-gene pairs within 1Mb of each other, there were 2,154,331 significant *cis*-eQTLs, and these were enriched for enhancer sequences in brain tissues, most strongly in DLFPC enhancers. There were also 45,453 significant *trans*-eQTLs, in which the SNP was more than 1Mb from the gene whose expression it correlated with. Only pair-wise comparisons of SNPs and genes were carried out and there was no report of whether the expression of any genes correlated with the PRS, although it was reported that the PRS was higher in cases than controls.

If multiple common variants exert their effect indirectly through regulatory networks, it is plausible that the PRS might reflect a cumulative effect on the expression of one or more "core" genes. This cumulative effect might be greater than any pairwise effect due to a single SNP. Also, it might be more pronounced for core genes than for the genes which only exerted their effect indirectly and so could serve to focus attention on core genes. Hence, it seems logical to explore whether the schizophrenia PRS is associated with gene expression.

## Materials and methods

The dataset used in the previous gene expression study was downloaded from the CommonMind Consortium (CMC) Knowledge Portal (*https://www.synapse.org/#!Synapse:syn2759792/wiki/69613*) consisting of SNP genotypes and RNAseq results from DLPFC samples originating from tissue collections at Mount Sinai NIH Brain Bank and Tissue Repository (MSSM), University of Pennsylvania Brain Bank of Psychiatric illnesses and Alzheimer’s Disease Core Center (Penn) and The University of Pittsburgh NIH NeuroBioBank Brain and Tissue Repository (Pitt), collectively referred to as the CMC MSSM-Pitt-Penn dataset (Fromer et al., 2016).

Genotypes and expression levels were available for 258 subjects with schizophrenia and 279 controls. The distributions of ethnicities were reported to be similar between subjects with schizophrenia and controls (Caucasian 80.7%, African-American 14.7%, Hispanic 7.7%, East Asian 0.6%). The methods for obtaining the genotypes and expression data have been described by the authors of the original study (Fromer et al., 2016). Genotyping was performed on the Illumina Infinium HumanOmniExpressExome 8 v 1.1b chip (Catalog #: WG-351-2301) using the manufacturer’s protocol. QC was performed using PLINK to remove markers with: zero alternate alleles, genotyping call rate < 0.98, Hardy-Weinberg p-value < 5 × 10^−5^, and individuals with genotyping call rate < 0.90. Marker alleles were phased to the forward strand, and ambiguously stranded markers have been removed. The gene expression data had been obtained by RNA sequencing of tissue from the DLPFC followed by adjustment for ancestry and other appropriate covariates.

In order to obtain a polygenic risk score (PRS) for schizophrenia, the file called *scz2.prs.txt.gz*, containing ORs and p values for 102,636 SNPs, was downloaded from the Psychiatric Genetics Consortium (PGC) website (*www.med.unc.edu/pgc/results-and-downloads*). This training set was produced as part of the previously reported PGC2 schizophrenia GWAS (Schizophrenia Working Group of the Psychiatric Genomics Consortium, 2014). SNPs with p value < 0.05 were selected and their log(OR) summed over sample genotypes using the --*score* function of *plink 1.09beta* in order to produce a PRS for each subject (www.cog-genomics.org/plink/1.9/) (Chang et al., 2015; Purcell et al., 2007, 2009). The first 20 principal components for all genotyped SNPs were produced using the -- *pca* and --*make-rel* functions of *plink*.

Statistical tests and data manipulation were carried out using R version 3.3.2 (R Core Team, 2014).

The PRS was higher in cases than controls (t = 3.5, p = 0.00059) but it was noticed that the association became much more highly significant when the SNP principal components were included as covariates (p = 2.5*10^−17^). This seemed to be due the fact that the first principal component was very strongly negatively correlated with the PRS (r = −0.81, p = 1.2*10^−126^). This correlation was present in both the controls (r = 0.85, p = 1.8*10^−78^) and the cases (r = −0.80, p = 2.5*10^−59^). The plots of controls and cases of the first and second principal components are shown in Figures 1A and 1B. It can be seen that the distribution is similar in both groups. Most subjects have a very high value for the first principal component and a very low value for the second one but several subjects are clearly outliers from this main grouping and have low scores for the first principal component or, in a few cases, high scores for the second. In order to filter out these outliers, the dataset was narrowed to include subjects with the first principal component value greater than 0.01 and second principal component value less than 0.04. This left 207 controls and 209 cases and produced a more homogeneous sample in which the PRS was not significantly correlated with the first principal component but was very significantly higher in cases than controls (t = 8.3, p = 1.3*10^−15^).

**Figure 1.**
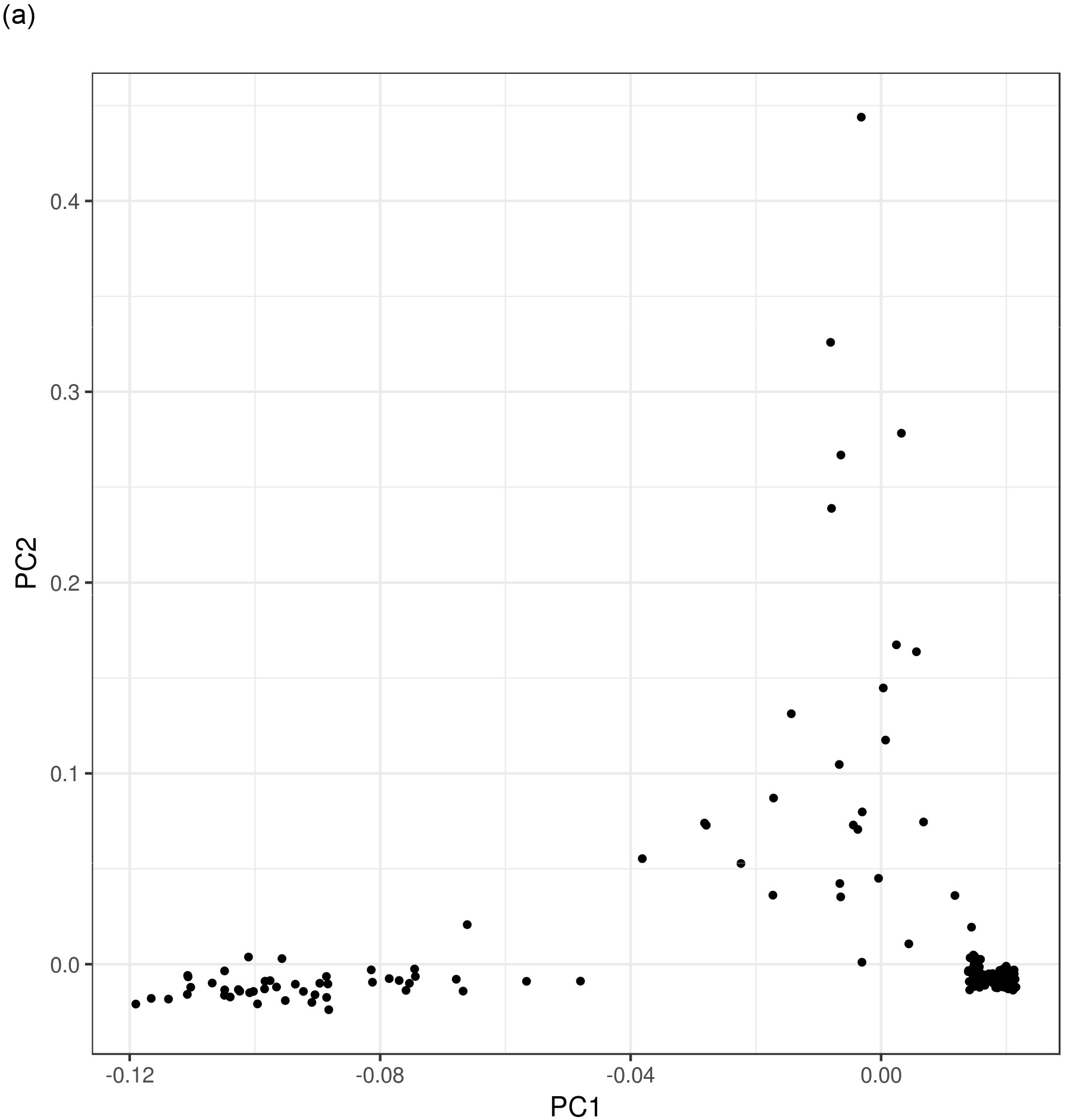
Controls (a) and subjects with schizophrenia (b) plotted against first two principal components from SNP genotypes.

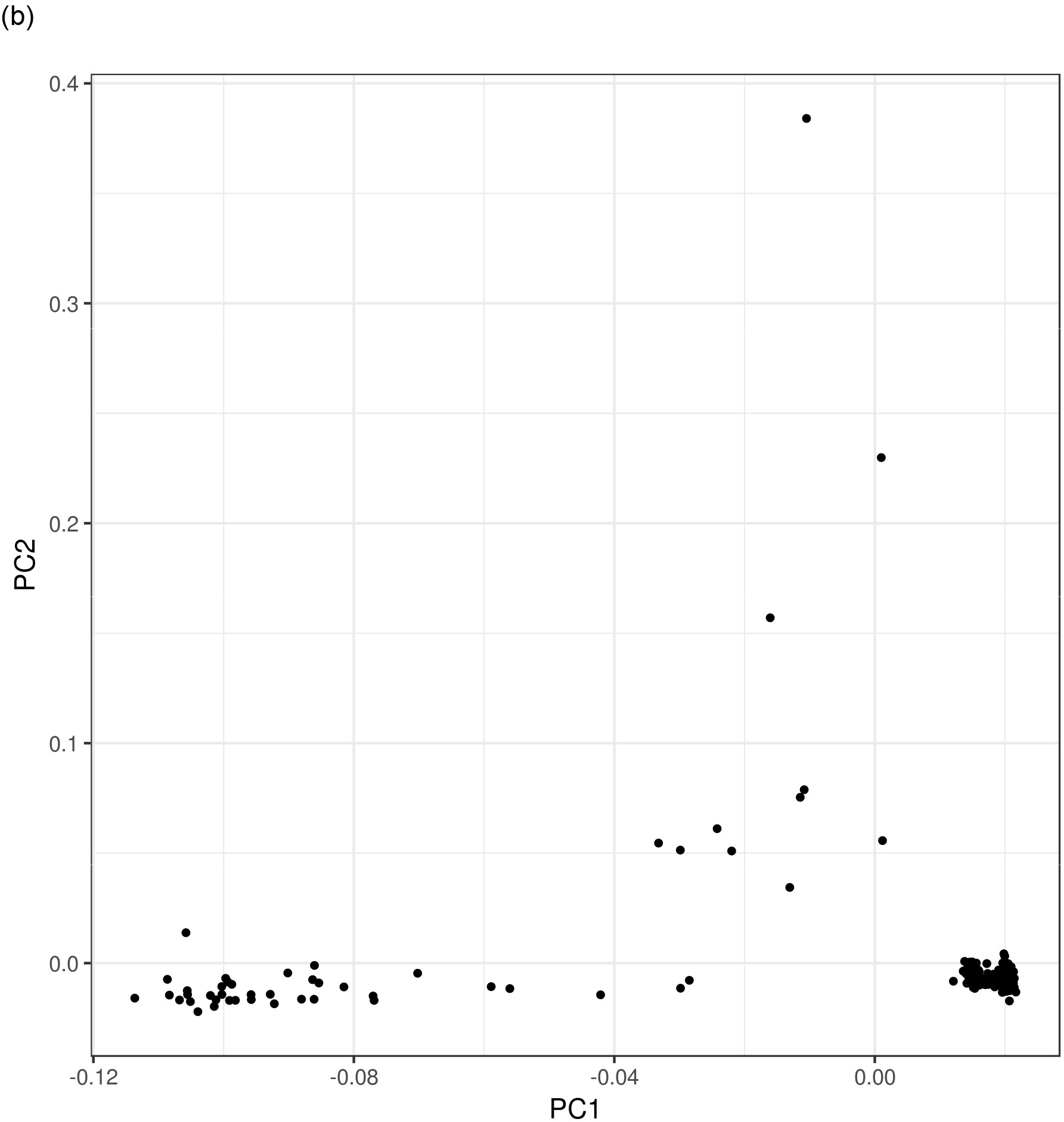

Regression of the expression level of each of 16,423 genes was performed on the PRS including the 20 principal components as covariates. The minus log base 10 of the p value (MLP) was recorded for each gene, as was the signed log p value (SLP), which is the same as the MLP but given a positive sign if the regression coefficient is positive and a negative sign if the coefficient is negative.

As well as the analyses based on individual genes, gene set analyses were performed to see if it was possible to identify a set of genes whose expression tended to correlate with the PRS. Two lists of gene sets were used. The first consisted of the 32 gene sets reported to be enriched for ultra-rare, gene disruptive variants in a sample of exome-sequenced schizophrenia cases (Genovese et al., 2016). These included categories such as genes expressed in neurons, genes near GWAS hits and genes which are loss of function intolerant. The second group consisted of 1454 "all GO gene sets, gene symbols" pathways downloaded from the Molecular Signatures Database at http://www.broadinstitute.org/gsea/msigdb/collections.jsp (Subramanian et al., 2005).

It is challenging to detect whether the expression pattern of a set of genes correlates with the PRS. One cannot simply test whether genes in the set are more strongly correlated with PRS than those not in the set because it is expected that the expression patterns of genes within a set will correlate with each other so that their correlations with PRS are not expected to be independent. Nor can one test whether the mean or total expression of members of the set correlates with PRS because it is expected that if a biological effect is present then expression of some genes will be increased and others decreased. Instead, the MLPs for each gene in the set were summed and the total compared with that which would be expected by chance by carrying out permutations of the PRS across subjects and then recalculating and totalling the gene-wise MLPs for each permuted dataset. The proportion of times the totalled MLPs for permuted datasets exceeded the total for the real dataset was used to obtain an empirical p value, and minus log base 10 of this empirical p value was used to obtain an MLP for the gene set.

## Results

The Q:Q plot for the SLPs for the correlation of PRS with the expression of each of 16,423 genes are shown in Figure 2. It can be seen that the results almost exactly follow what would be expected under the null hypothesis if the expression patterns of the genes were independent. The most highly significant correlation was for SULF1 with p=0.000005 (MLP=5.3) but this would not reach conventional standards of significance if a Bonferroni correction were applied. SULF1 codes for an extracellular heparan sulfate endosulfatase and does not seem to be a likely candidate for involvement in the aetiology of schizophrenia. It could be argued that expression patterns of genes are likely to be correlated with each other and hence that a Bonferroni correction might be overly conservative. Table 1 lists the genes achieving MLP>3.

**Figure 2.**
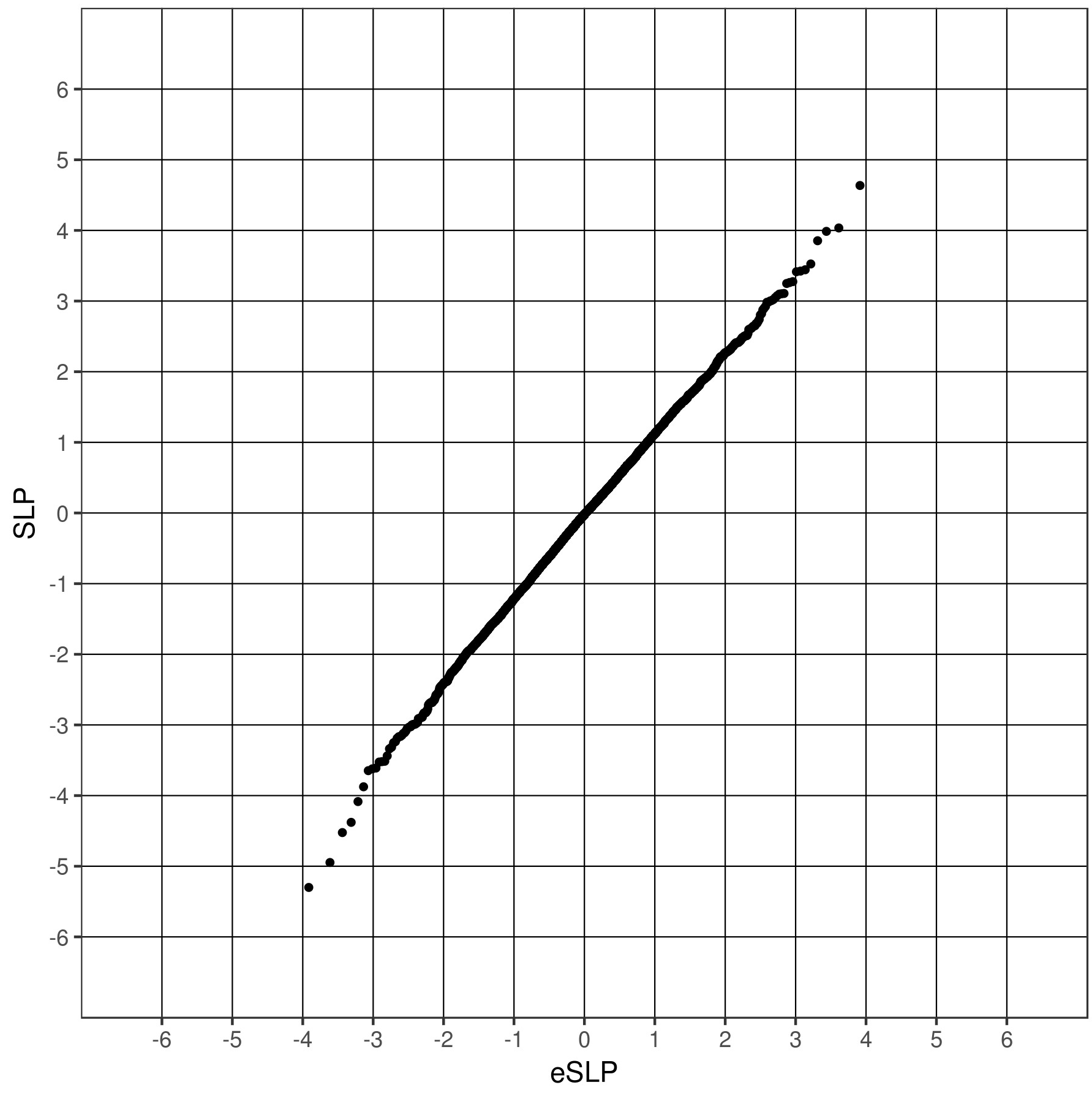
Q:Q plot of observed versus expected values for gene-wise SLPs assessing correlation between PRS and gene expression.

**Table 1.**
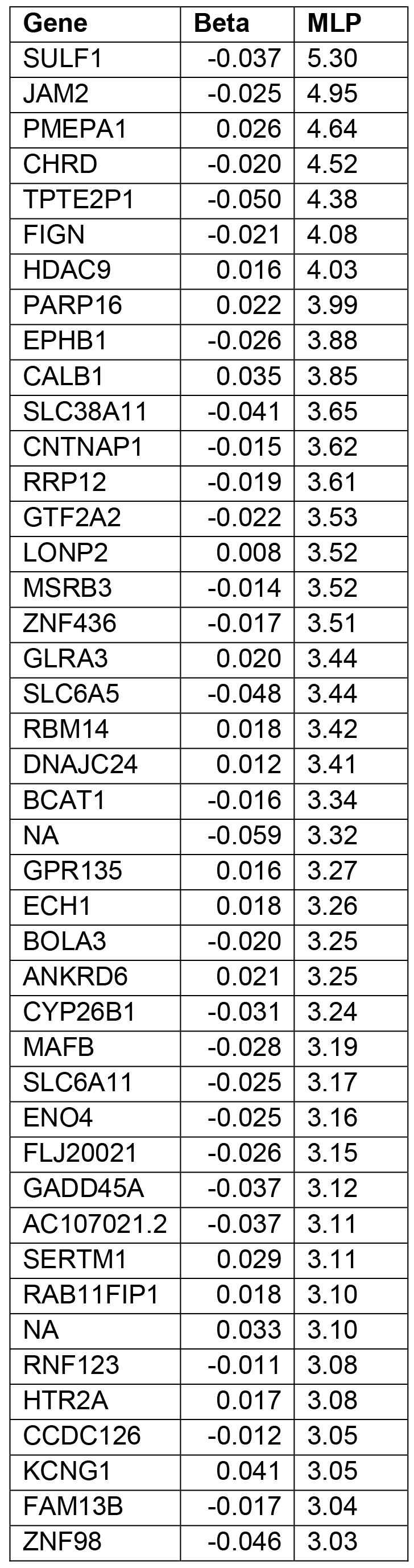
List of genes for which the correlation of PRS with expression is significant at p<0.001 (MLP>3).

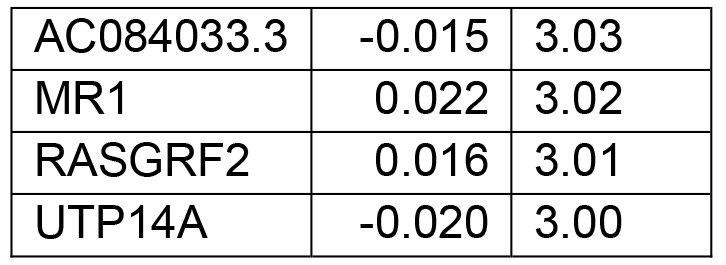

Table 2 shows the permutation-derived MLPs for each of the gene sets previously used to test for enrichment for damaging ultra-rare variants. It can be seen that none of the sets produces highly significant evidence for correlation with PRS. For only two of the sets, *alid* and *mir137*, does the MLP exceed 2 and again these results would not be significant if a Bonferroni correction were applied. Again, it could be argued that this might be conservative because the sets overlap each other and hence the MLPs, although individually valid, are not independent.

**Table 2.** Permutation-derived MLPs for each of the 32 gene sets previously used to test for enrichment for damaging ultra-rare variants in schizophrenia. The definition of each set is provided in the online methods section of the original publication (Genovese et al., 2016).

| Gene set | Symbol | MLP |
| --- | --- | --- |
| OMIM intellectual disability (Hamosh et al., 2005) | *alid* | 2.22 |
| Expression specific to brain (Fagerberg et al., 2014) | *brain* | 0.00 |
| Bound by CELF4 (Wagnon et al., 2012) | *celf4* | 0.18 |
| Missense-constrained (Samocha et al., 2014) | *constrained* | 0.34 |
| Involved in developmental disorder (Deciphering Developmental Disorders Study, 2017) | *dd* | 0.34 |
| De novo variants in autism (Fromer et al., 2014) | *denovo.aut* | 1.05 |
| De novo variants in coronary heart disease (Fromer et al., 2014) | *denovo.chd* | 0.08 |
| De novo variants in epilepsy (Fromer et al., 2014) | *denovo.epi* | 0.34 |
| De novo duplications in ASD (Kirov et al., 2012) | *denovo.gain.asd* | 0.38 |
| De novo duplications in bipolar disorder (Kirov et al., 2012) | *denovo.gain.bd* | 0.11 |
| De novo duplications in schizophrenia (Kirov et al., 2012) | *denovo.gain.scz* | 0.65 |
| De novo variants in intellectual disability (Fromer et al., 2014) | *denovo.id* | 0.18 |
| De novo deletions in ASD (Kirov et al., 2012) | *denovo.loss.asd* | 1.04 |
| De novo deletions in bipolar disorder (Kirov et al., 2012) | *denovo.loss.bd* | 0.86 |
| De novo deletions in schizophrenia (Kirov et al., 2012) | *denovo.loss.scz* | 0.04 |
| De novo variants in schizophrenia (Fromer et al., 2014) | *denovo.scz* | 0.87 |
| Bound by FMRP (Darnell et al., 2011) | *fmrp* | 0.20 |
| Implicated by GWAS (Schizophrenia Working Group of the Psychiatric Genomics Consortium, 2014) | *gwas* | 0.92 |
| Targets of microRNA-137 (Robinson et al., 2015) | *mir137* | 2.17 |
| Expression specific to neurons (Cahoy et al., 2008) | *neurons* | 0.71 |
| NMDAR and ARC complexes (Kirov et al., 2012) | *nmdarc* | 0.04 |
| Loss-of-function intolerant (Lek et al., 2016) | *pLI09* | 0.41 |
| PSD-95 (Bayés et al., 2011) | *psd95* | 0.18 |
| Bound by RBFOX 1 or 3 (Weyn-Vanhentenryck et al., 2014) | *rbfox13* | 0.04 |
| Bound by RBFOX 2 (Weyn-Vanhentenryck et al., 2014) | *rbfox2* | 0.26 |
| Synaptic (Pirooznia et al., 2012) | *synaptome* | 0.15 |
| Escape X-inactivation (Cotton et al., 2013) | *x.escape* | 0.04 |
| X-linked intellectual disability, Genetic Services Laboratories of the University of Chicago (Géczy et al., 2009; Moeschler, 2008; Moeschler et al., 2006; Rauch et al., 2006) | *xlid.chicago* | 0.32 |
| X-linked intellectual disability, Greenwood Genetic Centre (Moeschler et al., 2006) | *xlid.gcc* | 0.36 |
| X-linked intellectual disability, OMIM (Hamosh et al., 2005) | *xlid.omim* | 0.32 |
| X-linked intellectual disability (combined) | *xlid* | 0.04 |

Of the 1454 GO gene sets, none produced a result which would be significant after Bonferroni correction. However 11 did produce MLPs exceeding 3 and these are listed in Table 3. Again, these are not independent but include overlapping sets of genes. In particular, the sets relating to histone modification contain RBM14, which has a gene-wise MLP of 3.42, whereas the regulation of cell differentiation sets all include MAFB, which has a gene-wise MLP of 3.19. Table 4 lists all the genes which occur in at least one of these sets and which have a gene-wise MLP>1.3, equivalent to p<0.05.

**Table 3.** GO gene sets achieving MLP>3 (out of 1454 sets in total).

| Gene set | MLP |
| --- | --- |
| CHROMOSOME_ORGANIZATION_AND_BIOGENESIS | 3.49 |
| ESTABLISHMENT_AND_OR_MAINTENANCE_OF_CHROMATIN_ARCHITECTURE | 3.48 |
| HISTONE_MODIFICATION | 3.44 |
| NEGATIVE_REGULATION_OF_CELL_DIFFERENTIATION | 3.42 |
| REGULATION_OF_MYELOID_CELL_DIFFERENTIATION | 3.41 |
| TRANSCRIPTION_FACTOR_COMPLEX | 3.36 |
| COVALENT_CHROMATIN_MODIFICATION | 3.35 |
| CHROMATIN_MODIFICATION | 3.31 |
| SENSORY_ORGAN_DEVELOPMENT | 3.18 |
| TRANSCRIPTION_COACTIVATOR_ACTIVITY | 3.12 |
| NEGATIVE_REGULATION_OF_MYELOID_CELL_DIFFERENTIATION | 3.07 |

**Table 4.** Genes from the GO sets with MLP>3 where the gene-wise MLP exceeds 1.3 (equivalent to p<0.05).

| Gene | beta | MLP |
| --- | --- | --- |
| HDAC9 | 0.016 | 4.03 |
| RBM14 | 0.018 | 3.42 |
| MAFB | -0.028 | 3.19 |
| ZMIZ2 | -0.015 | 2.83 |
| CHMP1A | 0.011 | 2.64 |
| JAG2 | -0.014 | 2.35 |
| FOXO3 | -0.008 | 2.22 |
| TAF11 | -0.011 | 2.14 |
| SNAPC4 | -0.014 | 2.05 |
| MAML2 | -0.013 | 1.89 |
| TGFB2 | -0.015 | 1.81 |
| SUPT4H1 | -0.008 | 1.76 |
| SMARCD3 | 0.010 | 1.70 |
| NSD1 | -0.005 | 1.67 |
| DKC1 | -0.009 | 1.65 |
| HDAC6 | 0.006 | 1.59 |
| HCFC1 | -0.008 | 1.55 |
| HUWE1 | -0.006 | 1.53 |
| SUPT16H | -0.007 | 1.51 |
| ERCC4 | -0.008 | 1.51 |
| CARTPT | 0.044 | 1.50 |
| HDAC3 | 0.008 | 1.48 |
| TERF2IP | -0.007 | 1.47 |
| ACIN1 | 0.006 | 1.41 |
| CENPE | 0.017 | 1.40 |
| GTF3C4 | -0.006 | 1.37 |
| PDS5B | -0.006 | 1.35 |
| SIRT5 | -0.007 | 1.32 |
| RNF14 | -0.006 | 1.32 |

## Discussion

This approach has failed to yield any genes or gene sets whose expression is strongly associated with the schizophrenia PRS. It would be possible to argue that the Bonferroni correction is overly conservative and that there may be some association but the main aim of the analysis, to provide a strong focus on a small set of "core" genes, has not been successful. The PRS represents a cumulation of genetic risk factors from multiple locations but it is not the case that the variants involved exert their effect on risk primarily by combining to effect the expression of a small number of genes. In terms of effects on gene expression, the analyses described here do not add anything to the results previously reported for pair-wise analyses (Fromer et al., 2016).

As an aside, it is worth noting the very strong correlation of the schizophrenia PRS with the first principal component of the ethnically heterogeneous sample. It has recently been noted that the PRS for schizophrenia stratifies by ancestry (Weiner et al., 2017). Likewise, the PRS for type 2 diabetes and coronary heart disease vary between populations and this must be taken account of if attempting to estimate an individual's risk (Reisberg et al., 2017). Although it may be expected that the mean PRS will vary somewhat according to ethnicity, it is perhaps a little surprising that the observed correlation with the first principal component is quite so strong and this will be the subject of further investigation. However this phenomenon does not seem to be relevant to the findings reported here, which were obtained from the more homogeneous sub-sample.

This study does not support the proposal that multiple common variants exert a joint effect on schizophrenia risk through a cumulative effect on the expression of a small number of genes. It is not hard to conceive of different mechanisms whereby the "polygenic risk" might be observed. One kind of mechanism is that a variant has some effect on gene function or expression which has a small but direct effect on the pathophysiological processes which can lead to schizophrenia. However, one can also conceive of indirect effects which could be very distant from the "core biology". There are number of environmental factors which appear to influence risk of schizophrenia, including obstetric complications, cannabis use, maternal viral infection, maternal malnutrition and childhood adversity (Cannon et al., 2002; Gage et al., 2017; Khandaker et al., 2013; Stilo et al., 2013; Xu et al., 2009). Any genetic variant which affects exposure to such risk factors will be expected to be associated with schizophrenia and these mechanisms may work either within the proband or within their parents. To give one example, as pointed out previously, smoking during pregnancy is a risk factor for the subsequent development of schizophrenia and the observed genetic association between smoking behaviour and schizophrenia might be mediated via this mechanism, so that a genetic predisposition to smoke in the mother is associated with increased risk of schizophrenia in her offspring (Curtis, 2017; Hartz et al., 2017; Niemel et al., 2016). Likewise, genetic variants which influence pelvis size, foetal head size, susceptibility to viral infection, cannabis preference and susceptibility to be either a victim or perpetrator of childhood abuse will all be expected to be associated with schizophrenia risk. Of course, such indirect effects may be very small but GWAS sample sizes are such that very small effects can be detected. If this is view is valid then a number of common variants may contribute to the "omnigenic" signal through indirect environmental mechanisms as well as through the interactive effects in intra- and inter- cellular networks which have been proposed (Boyle et al., 2017).

## Conflict of interest

The author declares he has no conflict of interest.

## Acknowledgments

The author thanks the CommonMind Consortium for making this dataset available. The CommonMind Consortium data generation was supported by funding from Takeda Pharmaceuticals Company Limited, F. Hoffman-La Roche Ltd and NIH grants R01MH085542, R01MH093725, P50MH066392, P50MH080405, R01MH097276, RO1-MH-075916, P50M096891, P50MH084053S1, R37MH057881 and R37MH057881S1, HHSN271201300031C, AG02219, AG05138 and MH06692. Brain tissue for the study was obtained from the following brain bank collections: the Mount Sinai NIH Brain and Tissue Repository, the University of Pennsylvania Alzheimers Disease Core Center, the University of Pittsburgh NeuroBioBank and Brain and Tissue Repositories and the NIMH Human Brain Collection Core. CMC Leadership: Pamela Sklar, Joseph Buxbaum (Icahn School of Medicine at Mount Sinai), Bernie Devlin, David Lewis (University of Pittsburgh), Raquel Gur, Chang-Gyu Hahn (University of Pennsylvania), Keisuke Hirai, Hiroyoshi Toyoshiba (Takeda Pharmaceuticals Company Limited), Enrico Domenici, Laurent Essioux (F. Hoffman-La Roche Ltd), Lara Mangravite, Mette Peters (Sage Bionetworks), Thomas Lehner, Barbara Lipska (NIMH).

